# A shear lag model of the podocyte foot process network predicts a mechanical feedback loop driving progressive effacement

**DOI:** 10.64898/2026.07.15.738799

**Authors:** Mingxuan Bi, Hanxun Jin, Pongpratch Puapatanakul, Yuxuan Huang, Chengqing Qu, Jeffrey H. Miner, Hani Y. Suleiman, Guy M. Genin

**Affiliations:** Department of Mechanical Engineering and Materials Science, Washington University in St. Louis, St. Louis, MO, USA; NSF Science and Technology Center for Engineering Mechanobiology; Department of Mechanical Engineering, University of Cincinnati, Cincinnati, OH, USA; Division of Nephrology, Department of Medicine, Washington University School of Medicine, St. Louis, MO, USA; Division of Nephrology and Department of Internal Medicine, UT Southwestern Medical Center, Dallas, TX, USA

**Keywords:** podocyte, shear lag, foot process, glomerular filtration, mechanobiology, FSGS

## Abstract

The podocyte foot process network forms the final barrier of the kidney’s glomerular filtration system. Under mechanical stress this network is prone to injury in which podocytes lose connectivity to their neighbors and begin the progression toward effacement, but what governs its mechanical resilience is unknown. We show that the network is built like a lap joint: two major processes coupled through interdigitating foot processes, a configuration that behaves as a classical shear lag system, with force concentrating at the joint ends and decaying over a characteristic transfer length set by geometry and stiffness. A discrete network model reproduces the continuum shear lag solution and identifies a hierarchy among governing parameters, with cytoskeletal stiffening of the major process amplifying foot process force more potently than basement membrane stiffness. Applying the model to morphometric data from puromycin aminonucleoside nephrosis, a model of human minimal change disease and early focal segmental glomerulosclerosis, reveals a mechanical positive feedback loop: force concentration drives foot process loss, which raises force on surviving segments and accelerates further loss. This nonlinear amplification implies a threshold beyond which failure becomes self-sustaining, analogous to the critical crack length in fracture mechanics.

## 1. Introduction

The kidney’s glomerular filtration barrier depends on podocytes, specialized epithelial cells whose interdigitating foot processes form the final mechanical barrier controlling passage of solutes from blood to urine. Failure of this barrier manifests as proteinuria and, when progressive, leads to focal segmental glomerulosclerosis (FSGS) and chronic kidney disease [1]. Although biochemical and genetic mechanisms of podocyte injury have been studied extensively [2–4], the filtration barrier is fundamentally a mechanical structure: foot processes are anchored to the glomerular basement membrane (GBM), subjected to hydrostatic and flow-induced forces, and must maintain structural integrity against varied mechanical loading throughout life [5]. Podocyte loss is largely irreversible in the adult kidney because these terminally differentiated cells have limited capacity for replacement [3, 6], making mechanical failure of the foot process network a key step in disease progression. During filtration, podocyte foot processes experience transmural pressure differences, which generate tensile stresses that distend the glomerular capillary and stretch the overlying podocyte network. Simultaneously, ultrafiltrate flow through the filtration slits produces shear drag on individual foot processes [5, 7]. Cytoskeletal prestress within the actin-rich foot process core adds mechanical loads [8, 9]. Experimental and computational studies establish that these forces actively drive podocyte behavior: mechanical overload from glomerular hypertension and hyperfiltration promotes podocyte detachment [1], and mechanosensitive ion channels such as TRPC6 transduce mechanical stimuli into biochemical signals that regulate foot process structure [10]. Foot processes display a non-random spatial organization on glomerular capillaries (Fig. 1). High-resolution 3D imaging shows that major and secondary foot processes adopt preferred orientations in healthy kidneys, that these orientations are established only after vascular maturation, and that they are lost in both nephrotoxic serum nephritis and podocin-mutant mice [11]. Foot process geometry is therefore mechanically meaningful and dynamically maintained, which raises the question of how the network distributes mechanical load and where structural failure is most likely to begin. The transfer of stresses across the interdigitating geometry of the podocyte foot process network resembles the shear lag phenomenon in composite materials and adhesive joints. Shear lag theory describes how axial load in adherends is transferred via shear through a compliant interfacial layer, producing characteristic force gradients that concentrate stress near the ends of the overlap region [12, 13]. In the podocyte, two major processes (the adherends) are mechanically coupled through interdigitating foot processes and the intervening GBM (the interfacial layer). This structural analogy suggests that classical shear lag mechanics may govern force transmission in the foot process network. Shear lag approaches are attractive because they are analytically tractable and yield scaling laws with testable predictions. Experimental measurements of foot process forces remain technically challenging and are typically limited to aggregate or averaged quantities [14]. A shear lag framework yields analytical solutions with well-understood scaling laws expressed in terms of physical parameters such as process geometry, membrane stiffness, and intraglomerular pressure. We developed a shear lag model for the podocyte foot process network based upon a discrete representation of interdigitating foot processes. After verifying it against the classical analytical shear lag solution, we applied it to characterize how major process stiffness, membrane stiffness, and foot process orientation govern the magnitude and spatial distribution of forces across the network. To test whether the model can explain spatial patterns of injury observed in disease, we applied it to published morphometric data from puromycin aminonucleoside (PAN) nephrosis in 6 week-old male Wistar rats [15]. PAN nephrosis produces progressive foot process effacement that recapitulates key features of human minimal change disease and early FSGS [16–18]. Miyaki et al. [15] documented systematic, spatially patterned structural changes following PAN treatment using serial block-face scanning electron microscopy: in healthy controls, foot processes were approximately perpendicular to the major process and distributed symmetrically along its length; by Day 2 post-injection, the total number of foot processes had decreased and processes near the ends of the major process had adopted lower angles, while central processes retained near-normal geometry; by Day 3, processes at the major process ends had disappeared entirely, while some central foot processes persisted. This spatially ordered progression, with edge loss preceding central loss, is difficult to explain by biochemical injury alone, because a diffusible toxin such as PAN would be expected to affect all foot processes roughly equally, and suggests a mechanical contribution. We hypothesized that shear lag mechanics governs force transmission between adjacent foot processes, that the resulting force concentrations explain the spatial pattern of injury documented by Miyaki et al., and that disease-driven loss of foot processes creates a mechanical positive feedback loop of progressive effacement.

**Figure 1.**
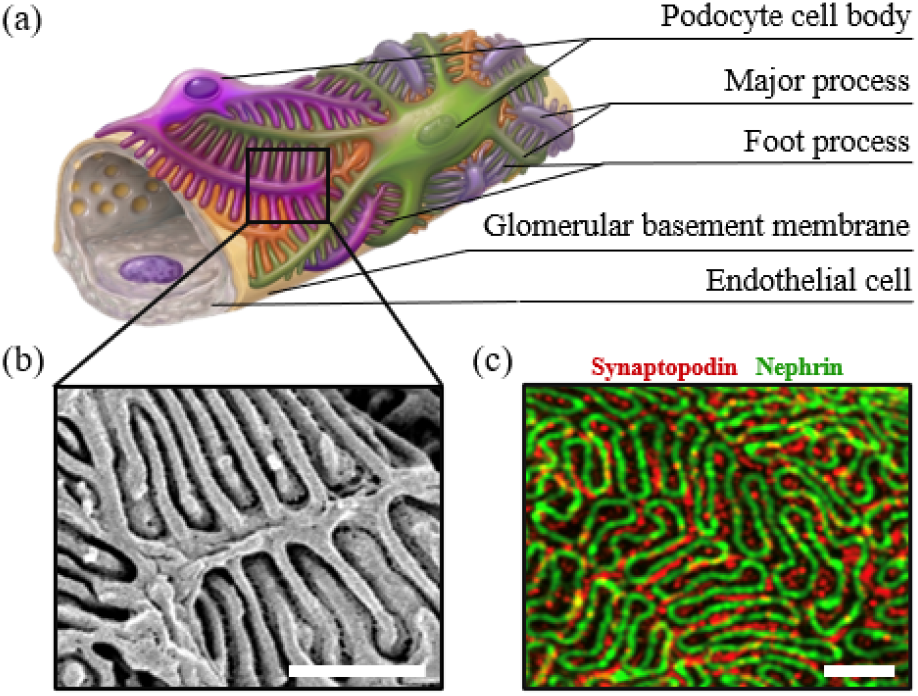
The podocyte foot process network of the glomerular filtration barrier. (*a*) Schematic of a podocyte wrapping a glomerular capillary, showing the hierarchical organization of cell body, major processes, and interdigitating foot processes anchored to the underlying glomerular basement membrane (made using BioRender.com). (*b*) Scanning electron micrograph of the filtration barrier in cross-section, resolving the foot processes, the slit diaphragms that bridge them, and the underlying GBM. Scale bar: 1 *µ*m. (*c*) Super-resolution micrograph of the apical filtration surface, showing the dense interdigitating foot-process network through which ultrafiltrate passes. Scale bar: 4 *µ*m.

## 2. Methods

### 2.1. Animal Studies

This study used previously published morphometric data, obtained from figures in Miyaki et al. (2020); no new animal or human experimentation was performed for this, and no ethical approval was required for the modelling and data-reuse components. For the representative imaging in Fig. 1, animal procedures were approved by the Institutional Animal Care and Use Committee of Washington University in St. Louis under protocol number 24-0095 and conducted in accordance with NIH Guide for the Care and Use of Laboratory Animals.

### 2.2. Electron Microscopy

Glomeruli were isolated using a Dynabeads-assisted glomerular isolation method as previously described [19]. Isolated glomeruli were fixed overnight in phosphate buffered saline containing 4% paraformaldehyde and 2% glutaraldehyde. Samples were rinsed in 0.15 M sodium cacodylate buffer and subsequently post-fixed with 1% osmium tetroxide prepared in 0.15 M sodium cacodylate buffer supplemented with 2 mM CaCl_2_ (pH 7.4) for 1 h at room temperature in the dark. Following three 10-min washes in distilled H_2_O, samples were dehydrated through an ascending ethanol series (50%, 70%, 90%, 100%, and 100%) for 10 min each. Dehydrated specimens were critical point dried using a Leica CPD300 system, mounted onto aluminum stubs with conductive carbon adhesive tabs, and coated with a 6 nm iridium layer using a Leica ACE600 sputter coater. Imaging was carried out on a Zeiss Merlin scanning electron microscope equipped with a Gemini II electron column operating at 3 keV and 200 nA. Secondary electron signals were collected using an Everhart-Thornley SE2 detector.

### 2.3. Ultrastructure Expansion Microscopy and Airyscan Imaging

Ultrastructure expansion microscopy was performed according to a previously established kidney tissue protocol [20]. Briefly, paraformaldehyde-perfused mouse kidney sections were incubated with anchoring reagents to enable covalent linkage of proteins to an acrylamide-based swellable hydrogel. Tissue embedding was achieved through in situ gel polymerization. Following polymerization, gel-embedded samples underwent denaturation in buffer containing 200 mM SDS, 200 mM NaCl, and 50 mM Tris-HCl (pH 9) at 95° C for 1.5 h. This treatment facilitated uniform tissue expansion while maintaining overall structural integrity. Gels were then expanded in distilled water to achieve an approximate fourfold linear expansion. Expanded tissues were immunostained with antibodies against renal structural proteins including nephrin and synaptopodin. Imaging was performed using a Zeiss Airyscan confocal microscope with a 60*×* objective lens. Acquisition settings included transmitted light (TD) imaging together with 405 nm, 488 nm, and 647 nm laser channels to evaluate tissue morphology and fluorescence labeling. Z-stack image series were acquired to visualize three-dimensional glomerular structures, and maximum intensity projections were generated for analysis. Images were deconvoluted and processed with uniform brightness and contrast adjustments applied consistently across all samples.

### 2.4. Shear Lag Model for a Foot Process Unit

We developed a discrete shear lag model of force transmission through an interdigitating podocyte foot process network (Fig. 2). The model geometry comprises two parallel major processes (adherends) coupled by an array of oblique foot processes that bridge the glomerular basement membrane (GBM), forming a configuration analogous to a lap joint in composite mechanics. Each foot process transmits force between a major process and the GBM through several mechanically distinct regions: a connection zone at the major process junction, the structural core of the foot process itself, and, for processes that contact the GBM, the intervening membrane. We represented each of these regions as a linear element arranged in series (Fig. 2). The effective stiffness of a lower foot process, which spans from the major process to the GBM, is therefore

**Figure 2.**
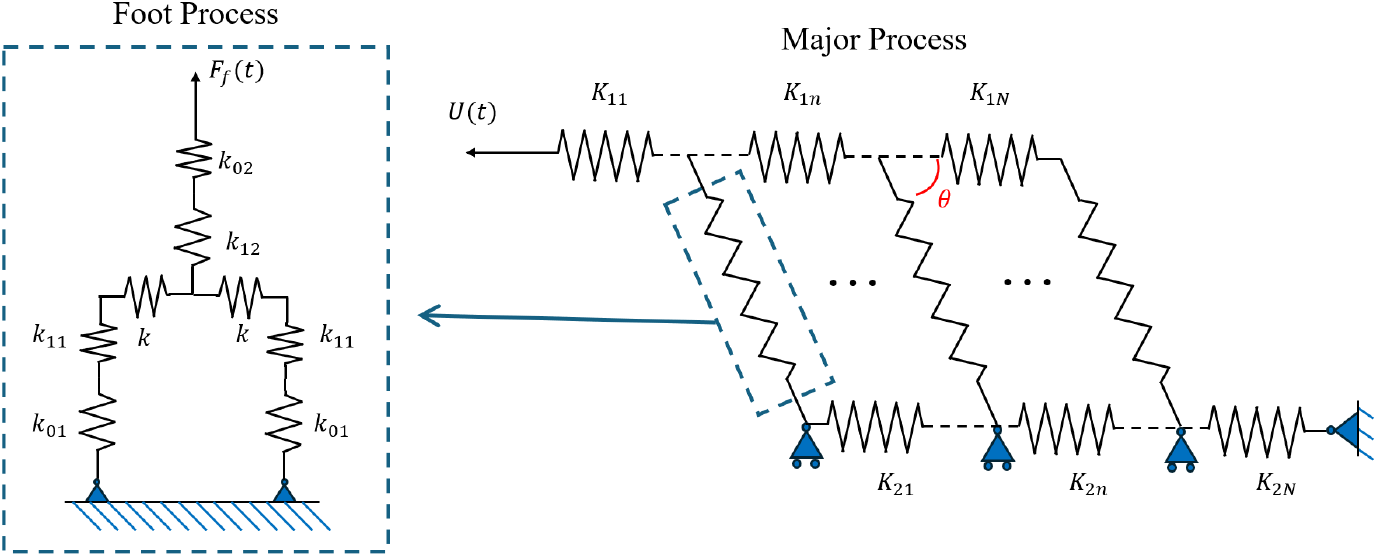
Schematic of the foot process network model illustrating the shear lag configuration. Major processes (horizontal) are connected by interdigitating foot processes (diagonal) modelled as elastic springs.

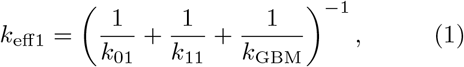

and the effective stiffness of an upper foot process, which does not contact the GBM, is

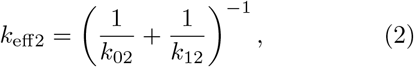

where *k*_01_ and *k*_02_ are the connection stiffnesses where each foot process joins its major process, *k*_11_ and *k*_12_ are the structural stiffnesses of the lower and upper foot process cores, and *k*_GBM_ is the stiffness of the GBM between adjacent attachment sites. At each node on the GBM, two lower foot processes converge from opposite major processes and act in parallel. To ensure that their combined stiffness equals that of a single upper foot process at the same node, each lower element was assigned half the stiffness of the corresponding upper element: *k*_01_ = *k*_02_*/*2 and *k*_11_ = *k*_12_*/*2. The GBM itself enters the model as the process-to-process connection *k*_GBM_ (written *k* in the parametric studies); biologically, this element lumps the compliance of the GBM with that of the integrin (*α*_3_*β*_1_)- and dystroglycan-mediated adhesions that anchor each foot process to it. A constant displacement Δ was imposed across each foot process unit, representing the strain imposed on the foot process network by capillary distension under transmural pressure. For a single unit, the elongations of the lower and upper foot processes were:

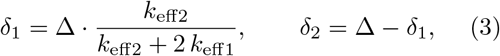

and the force transmitted through the unit was

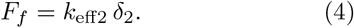

### 2.5. Assembly of the Foot Process Network

The single-unit formulation was extended to a full network in which two major processes are coupled through *N* interdigitating foot process segments. Each major process was discretized into segments and assembled into a global stiffness matrix following standard procedures for discrete elastic networks [21]:

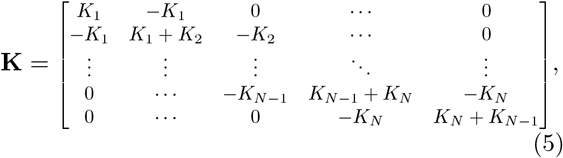

where *K*_*n*_ = *E*_*T*_ *hw/L*_*n*_ is the axial stiffness of the *n*th major process segment, with *E*_*T*_ the elastic modulus, *h* and *w* the cross-sectional height and width, and *L*_*n*_ the segment length. Nodal equilibrium gives:

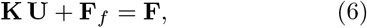

where **U** is the nodal displacement vector, **F**_*f*_ is the vector of foot process forces acting on the major process nodes, and **F** is the external force vector. Because foot processes are inclined at an angle *θ* to the major process axis, axial displacement of the major processes changes both the length and the orientation of each foot process. To capture this coupling, we computed the elongation of a foot process with rest length *l*_0_ from the deformed geometry:

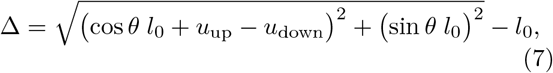

where *u*_up_ and *u*_down_ are the axial displacements of the upper and lower major process nodes to which the foot process is attached. The first term under the radical is the projection of the deformed foot process vector onto the major process axis; the second is the (unchanged) transverse projection. Because Δ depends nonlinearly on the nodal displacements, the system was solved iteratively.

### 2.6. Analytical Shear Lag Verification

To verify the discrete network model, we compared its predictions with a continuum shear lag solution derived by homogenizing the discrete connector stiffnesses into an equivalent interfacial shear modulus. The combined connector stiffness is

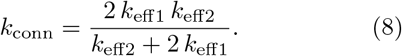

The mapping from discrete connectors to a continuum interfacial layer follows a simple homogenization. Equating the force per connector, *k*_conn_*δ*, to the shear traction integrated over the tributary area *L*_sec_ *× w* yields the equivalent shear modulus *G* = *k*_conn_ *H/*(*L*_sec_ *w*), where *H* = *l*_0_ is the interfacial layer thickness. For two adherends with moduli and thicknesses {*E*_1_, *t*_1_} and {*E*_2_, *t*_2_}, the governing shear lag parameter is

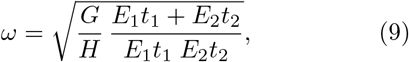

and the analytical shear stress distribution along the overlap length *L*_*T*_ is then

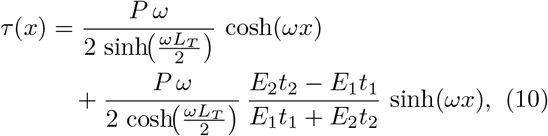

where *x* is measured from the midpoint of the overlap region and *P* = *F*_*f*,max_*/w* is the axial load per unit width, with *F*_*f*,max_ taken from the discrete calculation. For the case of equal adherends (*E*_1_ *≈ E*_2_ and *t*_1_ *≈ t*_2_), these reduce to

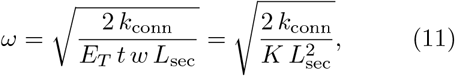

where *K* = *E*_*T*_ *hw/L*_sec_ and *t* = *h*, and

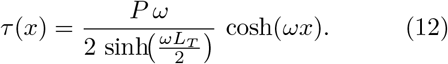

The shear force at each node was obtained by multiplying *τ* (*x*) by the foot process cross-sectional area.

Both discrete and analytical profiles were normalized by their respective maxima so that comparisons isolate spatial distributions.

### 2.7. Parametric Studies

Table 1 summarizes the key model parameters and their descriptions. The sensitivity of the force distribution to model parameters was examined through three sets of single-parameter sweeps. First, the ratio of GBM stiffness to foot process stiffness, *k/k*_eff2_, which controls the degree of force non-uniformity along the major process and reflects changes in basement membrane composition. Second, the major process stiffness ratio *K/K*_base_, which governs the overall force magnitude and represents cytoskeletal remodelling of the major process. Third, the initial foot process angle *θ* relative to the major process axis, which determines geometric coupling between axial and transverse deformations and varies with foot process morphology. In each sweep, the parameter of interest was varied over its full range while all other parameters were held at baseline values. A constant total displacement of Δ = 0.02 *l*_0_ was applied in all cases, corresponding to a 2% engineering strain of the foot process rest length.

**Table 1.** Model parameters and their descriptions.

| Variable | Description | Units |
| --- | --- | --- |
| $k_{11}, k_{12}$ | Structural stiffness of lower and upper foot processes | $\text{N m}^{-1}$ |
| $k_{01}, k_{02}$ | Connection between major process and foot process | $\text{N m}^{-1}$ |
| $k_{\text{GBM}}$ | Glomerular basement membrane stiffness | $\text{N m}^{-1}$ |
| $k_{\text{eff1}}, k_{\text{eff2}}$ | Effective stiffnesses of lower and upper foot process assemblies | $\text{N m}^{-1}$ |
| $k_{\text{conn}}$ | Combined connector stiffness used in continuum homogenization | $\text{N m}^{-1}$ |
| $K_n$ | Axial stiffness of the $n$-th major process segment | $\text{N m}^{-1}$ |
| $E_T$ | Elastic modulus of the major process | Pa |
| $h, w$ | Major process cross-sectional height and width | m |
| $L_T, L_n$ | Total and segment lengths of the major process | m |
| $F_f$ | Foot process force | N |
| $\delta_1, \delta_2$ | Elongation of lower and upper foot processes | m |
| $\Delta$ | Total imposed elongation ($0.02 l_0$) | m |
| $l_0$ | Foot process rest length | m |
| $\theta$ | Initial angle of foot process relative to major process axis | rad |
| $G$ | Effective shear modulus of the homogenized interfacial layer | Pa |
| $\omega$ | Shear lag parameter governing spatial decay of force | $\text{m}^{-1}$ |

### 2.8. Application to PAN Nephrosis

To examine how disease-driven structural changes alter force distribution, we applied the model to morphometric data from the literature for puromycin aminonucleoside (PAN) nephrosis in 6 week-old male Wistar rats [15]. PAN nephrosis produces progressive foot process effacement that recapitulates key features of human minimal change disease and early FSGS. Three morphometric quantities were extracted from published serial block-face scanning electron micrographs at each of three time points (healthy controls, Day 2 post-injection, and Day 3 post-injection): (i) foot process number *N*, (ii) foot process spacing along the major process, and (iii) angular orientation *θ* of each foot process relative to the major process axis. All three micrographs were analyzed at the same scale. For each image, the outlines of the major processes and the associated foot processes were independently traced, and the angular orientation between each foot process and the major process axis was measured. To reduce measurement bias, each angle was measured three times and the average value was used, and three independent measurement trials were conducted for each image. The resulting angles were used as inputs to the model to compute the force distribution at each disease stage and to evaluate how the force profile changes across disease progression. For each disease stage, the model was parameterized with the measured geometry while baseline stiffness values were held constant across conditions, so that differences in the predicted force profiles arise solely from the geometric changes associated with disease progression. The same constant displacement boundary condition (Δ = 0.02 *l*_0_) was applied in all cases.

### 2.9. Statistical Analysis

To assess whether predicted force distributions differed across disease stages, a scalar summary statistic was computed from each force profile: the area under the curve (AUC), obtained by trapezoidal integration of the foot process force over normalized axial position along the major process. Pairwise comparisons between disease stages were performed using Welch’s *t*-test with Holm correction for multiple comparisons. A one-way analysis of variance (ANOVA) was used to test for an overall effect of disease stage, with Tukey-Kramer post-hoc pairwise comparisons. A significance threshold of *p <* 0.05 was used throughout.

## 3. Results

### 3.1. Convergence of the Discrete Model to the Analytical Shear Lag Solution

The discrete spring network reproduced the characteristic force distribution predicted by classical shear lag theory: force was greatest at the ends of the overlap region, decayed toward the midpoint of the major process, and rose symmetrically beyond it (Fig. 3). This U-shaped profile is a hallmark of shear lag mechanics in composite lap joints and confirms that the interdigitating foot process architecture generates the same force concentration phenomenology. Quantitative agreement between the discrete and analytical models was close across the full normalized axial position when the number of foot process segments satisfied *N >* 5. For *N <* 5, the predictions of the discrete and continuous shear stress fields diverged, indicating a minimum segment density required for the continuum solution to be valid. For the foot process geometries considered here, *N* = 5 corresponds to roughly two segments per decay length 1*/ω*.

**Figure 3.**
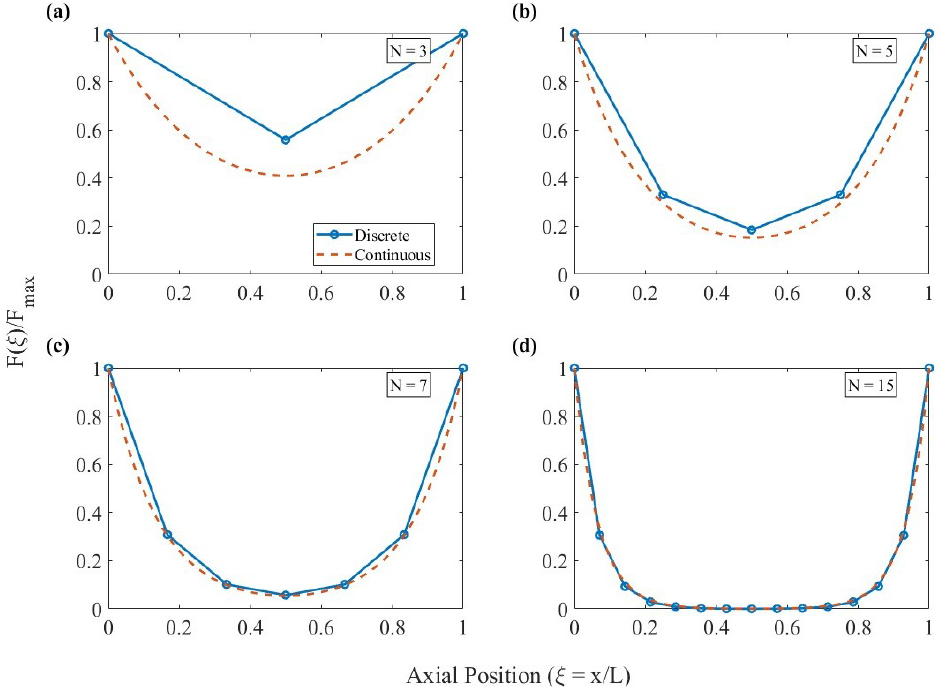
Convergence of the discrete spring network to the continuum shear lag solution. Normalised foot process force is plotted as a function of normalised axial position along the major process for the discrete model (markers) and the continuum solution (solid curve). Both profiles exhibit the characteristic shear lag pattern, with force maxima at the overlap ends and a minimum at the midpoint, and both agree closely for networks with more than five foot process segments (*N >* 5). Parameters: segment length *L*_sec_ = 2 *µ*m, foot-process length *l*_0_ = 3 *µ*m, major-process height and width *h* = *w* = 1 *µ*m, and angular orientation between major and foot processes *θ* = 90°. The axial segment stiffness is *K*_major_ = *E*_*T*_ *hw/L*_sec_. The corresponding stiffness ratios are *k*_eff2_*/k*_GBM_ = 0.5 and *K*_major_*/k*_GBM_ = 3.57 *×* 10^−3^.

### 3.2. Sensitivity to Stiffness Parameters

Two stiffness ratios govern the mechanical response of the foot process network. Each was varied independently while all other parameters were held at their baseline values.

#### GBM-to-foot-process stiffness ratio, *k/k*_eff2_

Increasing *k/k*_eff2_ produced progressively more nonuniform force distributions: forces concentrated at the ends of the major process while diminishing near its centre (Fig. 4a). This non-uniformity saturated at *k/k*_eff2_ *≈* 1; beyond this value, further increases in membrane stiffness had negligible additional effect on the force profile. Physically, saturation occurs because the foot process compliance, rather than the membrane compliance, becomes the limiting factor for force transmission once the membrane is sufficiently stiff.

**Figure 4.**
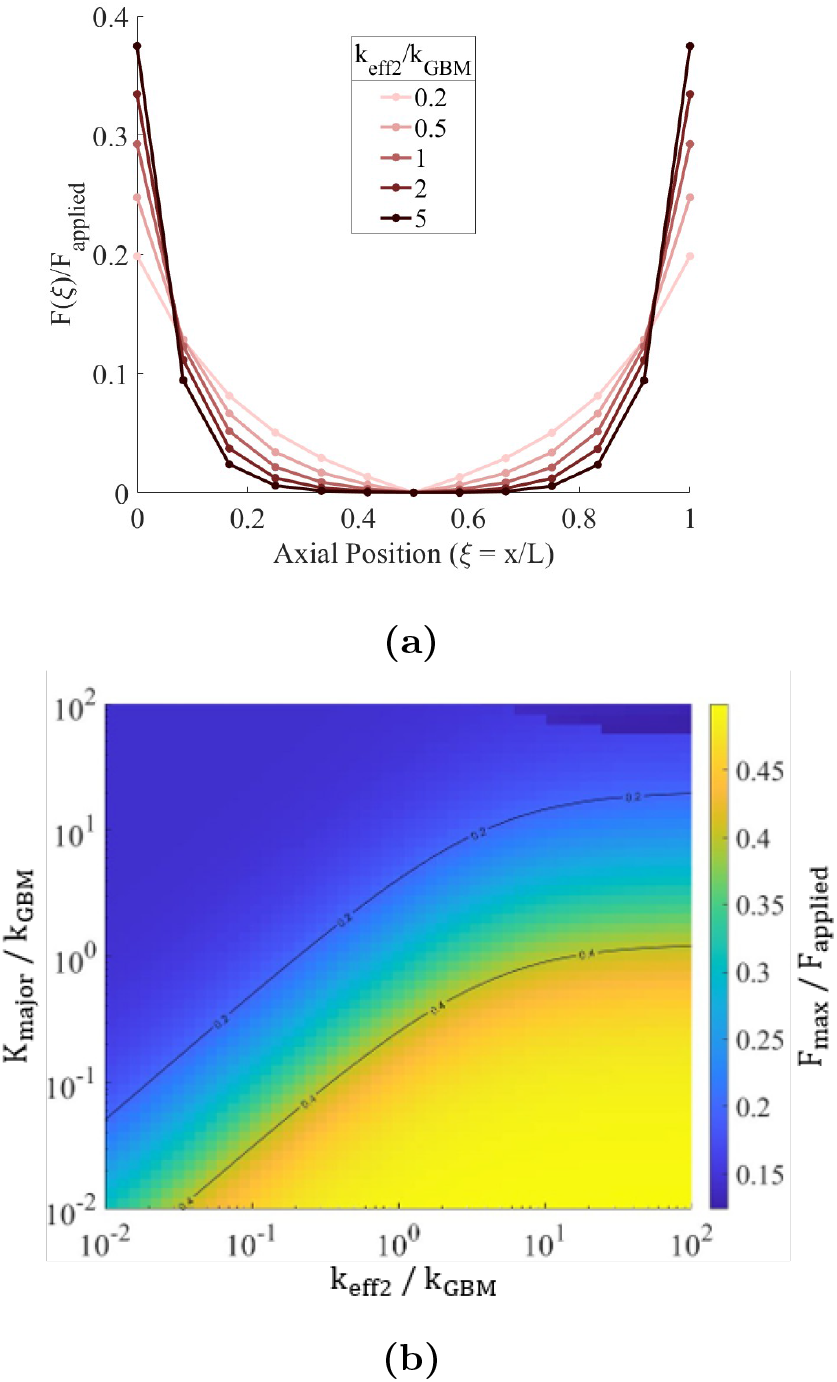
Effect of stiffness parameters on foot process force. (*a*) Normalised force along the major process for increasing ratios of GBM stiffness to foot process stiffness, *k/k*_eff2_, with *K/K*_base_ held constant. Force concentrates at segment ends and the distribution saturates near *k/k*_eff2_ *≈* 1. (*b*) Phase map of maximum foot process force as a function of both *K/K*_base_ (horizontal) and *k/k*_eff2_ (vertical), showing a low-force regime at *K/K*_base_ *<* 1 and a high-force regime in which major process stiffness dominates the response. Parameters: segment length *L*_sec_ = 2 *µ*m, foot-process length *l*_0_ = 3 *µ*m, major-process height and width *h* = *w* = 1 *µ*m, angular orientation between major and foot processes *θ* = 90°, and number of segments *N* = 13.

#### Major process stiffness ratio, *K/K*_base_

The phase map of maximum foot process force as a function of both stiffness ratios (Fig. 4b) revealed two distinct mechanical regimes. When *K/K*_base_ *<* 1, the major process is more compliant than the foot processes and deforms locally rather than transmitting load through the shear lag mechanism; foot process forces remain near zero regardless of membrane stiffness. When *K/K*_base_ *>* 1, the major process is stiff enough to engage the shear lag mechanism, and foot process force increases with both *K/K*_base_ and *k/k*_eff2_, with major process stiffness exerting the stronger influence. Quantitatively, when *k*_eff2_*/k*_GBM_ *<* 10, a tenfold increase in *K*_major_*/k*_GBM_ decreases the maximum normalized foot process force *F*_max_*/F*_applied_ by approximately 21% on average, whereas the same tenfold increase in *k*_eff2_*/k*_GBM_ increases it by about 29% on average. In this range, the largest increase produced by a tenfold increase in *k*_eff2_*/k*_GBM_ is 79.8%. When *k*_eff2_*/k*_GBM_ *>* 10, the influence of *k*_eff2_*/k*_GBM_ becomes much weaker (approximately 1% increase on average), while a tenfold increase in *K*_major_*/k*_GBM_ still produces an average reduction of roughly 27%. Here, the largest reduction produced by a tenfold increase in *K*_major_*/k*_GBM_ is 44.9%. This asymmetry has a direct biological implication: stiffening of the major process cytoskeleton, whether through actin bundling, intermediate filament upregulation, or crosslinker accumulation, amplifies mechanical loading on foot processes more than equivalent changes in GBM composition.

### 3.3. Sensitivity to Initial Foot Process Angle

The initial angle *θ* of the foot processes relative to the major process axis controls how axial displacement of the major processes projects into foot process elongation. The elongation depends on cos *θ* through the axial projection; at near-perpendicular orientations (*θ →* 90°), a given axial displacement produces a smaller change in the axial projection but a larger change in foot process length because the transverse component sin *θ l*_0_ dominates the deformed geometry. At shallow angles, the same displacement produces a smaller net change in foot process length, distributing forces more evenly and at lower magnitude. As *θ* increased toward perpendicular, both the maximum foot process force and the angular deformation at each segment increased, and their spatial distributions became more non-uniform (Fig. 5). The angular deformation profiles (Fig. 5a) show that segments near the ends of the major process undergo the largest changes in orientation, while central segments rotate relatively little. This spatial pattern mirrors the fore distribution (Fig. 5b) and identifies the end segments as the sites most susceptible to both mechanical overload and geometric remodelling. This prediction is tested against disease data in Section 3.4.

**Figure 5.**
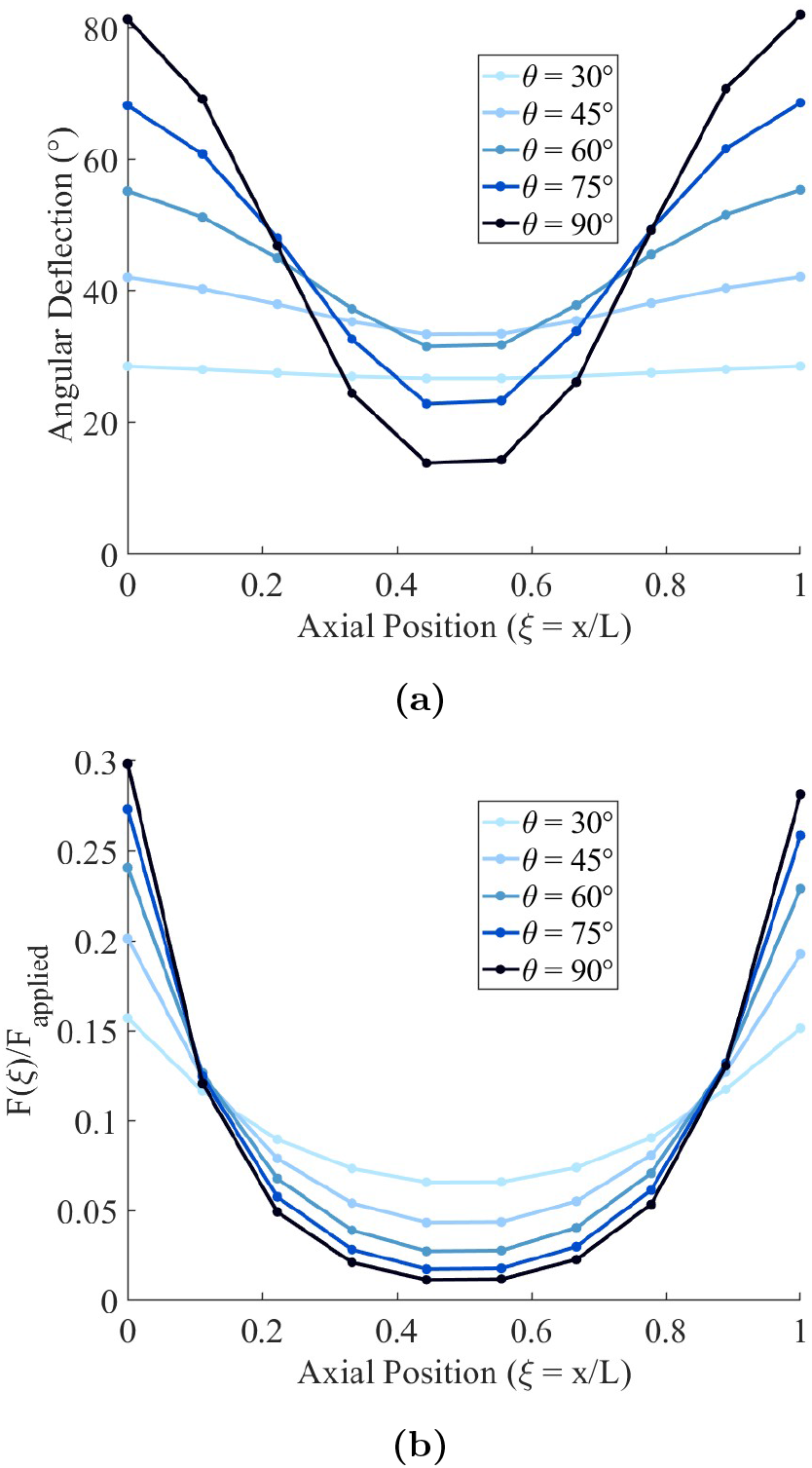
Effect of initial foot process angle *θ* on the mechanical response of the network. (*a*) Change in foot process angle at each segment along the major process for a range of initial angles. End segments undergo the largest angular deformation, consistent with the force concentration predicted by shear lag theory. (*b*) Normalised foot process force at each segment for the same set of initial angles. More perpendicular foot processes produce larger forces and steeper force gradients along the major process. Parameters: number of segments *N* = 10, and stiffness ratios *k*_eff2_*/k*_GBM_ = 0.4 and *K*_major_*/k*_GBM_ = 1.4.

### 3.4. Force Redistribution and Mechanical Vulnerability in Disease

Using the morphometric data from Miyaki et al. [15] described in Section 1 to parameterize the model at each disease stage, we applied the shear lag model under identical boundary conditions (Δ = 0.02 *l*_0_). The model revealed a progressive increase in both the magnitude and spatial non-uniformity of foot process forces (Fig. 6). As the number of foot processes decreased from healthy to Day 2 to Day 3, the force borne by each remaining segment increased: fewer structural elements must collectively resist the same imposed displacement, amplifying the force per element. The predicted force distributions differed significantly across all disease stages. Welch *t*-tests with Holm correction on scalar AUC values yielded *p* = 5.7 *×* 10^−5^ for Day 2 versus healthy and *p* =4.7 *×* 10^−6^ for Day 3 versus healthy, while Day 2 versus Day 3 also differed significantly (*p* = 7.1 *×* 10^−7^). A one-way ANOVA confirmed an overall effect of disease stage (*p* = 7.6 *×* 10^−11^), and Tukey-Kramer pairwise comparisons yielded adjusted *p <* 10^−6^ for all three pairs. The narrow, minimally overlapping 95% confidence intervals in Fig. 6 indicate that between-group differences substantially exceed within-group variability. Taken together, these results reveal a mechanical positive feedback loop. Force concentration at the ends of the major process promotes structural damage and foot process loss; loss reduces the number of load-bearing elements, which increases the force on each survivor; the elevated force drives further loss. The angular remodelling observed at Day 2, where edge processes adopt shallower angles before disappearing entirely by Day 3, is consistent with a transient geometric adaptation that partially offloads the most stressed elements but ultimately proves insufficient to arrest the cycle of progressive effacement.

**Figure 6.**
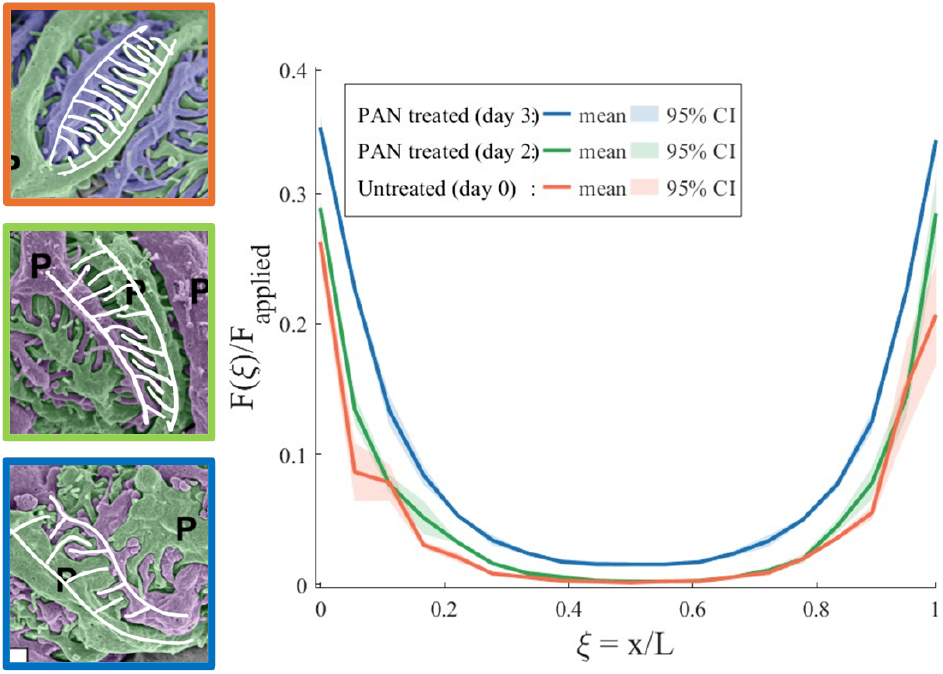
Progressive amplification of foot process force during PAN nephrosis. Predicted shear force is plotted as a function of normalised axial position along the major process for healthy controls (blue), Day 2 (orange), and Day 3 (red) post-injection in 6 week-old male Wistar rats. Solid curves show ean force profiles; shaded regions indicate 95% confidence intervals. Both the magnitude and spatial non-uniformity of force increase with disease progression, consistent with the reduction in foot process number and angular remodelling documented by Miyaki et al. [15]. All pairwise differences are statistically significant. For Day 2 vs untreated, Day 3 vs untreated, and Day 2 vs Day 3 respectively, Welch’s *t*-test with Holm correction gives *p* = 5.70 *×* 10^−5^, *p* = 4.68 *×* 10^−6^, and *p* = 7.13 *×* 10^−7^; Tukey-Kramer gives *p* = 3.81 *×* 10^−7^, *p* = 3.97 *×* 10^−19^, and *p* = 2.99 *×* 10^−14^.

## 4. Discussion

The shear lag framework developed here provides an analytical model for force distribution across the podocyte foot process network. The agreement between the discrete network and the continuum shear lag solution (Fig. 3) establishes that the interdigitating foot process architecture produces classical shear lag force concentration. The practical consequence is that the shear lag parameter *ω* (or, equivalently, the transfer length *λ* = 1*/ω*) provides a single, physically grounded metric that encodes foot process geometry, membrane stiffness, and major process modulus into one quantity governing the spatial decay of force. Because *ω* is expressed entirely in terms of measurable quantities, it enables direct comparison of mechanical states across conditions, genotypes, or species. The parametric analysis identifies a clear hierarchy among the governing parameters. Major process stiffness *K* controls whether the shear lag mechanism is active: below a threshold value, the major process deforms locally and does not transmit load to central foot processes. Above this threshold, *K* amplifies foot process forces more potently than membrane stiffness *k*, while force nonuniformity saturates once *k/k*_eff2_ *≈* 1. This asymmetry predicts that cytoskeletal stiffening of the major process through actin bundling proteins is a more consequential driver of foot process overload than changes in GBM composition, and that therapeutic strategies targeting GBM stiffness face diminishing returns beyond a critical stiffness. Foot process angle *θ* governs the severity of end-concentrated forces through its control of the geometric projection between axial displacement and foot process elongation. The model predicts that angular deformation and force are both greatest at the ends of the major process, identifying these as the sites most vulnerable to mechanical failure. Reducing *θ* decreases local force but, because transmural pressure is set by systemic haemodynamics, necessarily widens the filtration slits and compromises barrier selectivity. The mechanical trade-off of force reduction at the cost of barrier integrity is consistent with the clinical observation that proteinuria often precedes overt structural injury [1], and suggests that proteinuria onset may mark the point at which geometric adaptation begins to compromise the slit diaphragm. It is also consistent with the disease-associated loss of preferred foot process orientation reported by Unnersjö-Jess et al. [11], in which the orderly geometric arrangement of foot processes seen in healthy mice becomes randomized in both immune-mediated and genetic glomerular disease. Within our framework, geometric randomization distributes the imposed displacement across foot processes of more variable axial projection cos *θ*, disrupting any mechanical tuning present in the healthy configuration. The local remodelling our model predicts at the ends of the major process and the global loss of orientation observed by Unnersjö-Jess et al. may therefore be two manifestations of the same underlying mechanical adaptation, operating at different scales of the network. These two findings suggest the existence of a positive mechanical feedback loop. Force concentration at segment ends promotes foot process loss; loss increases the force on survivors; elevated force drives further loss. Because the shear lag decay length *λ* is a material property that does not change as segments are lost, removing end segments shifts the force maxima inward to the new terminal segments, producing nonlinear force amplification with decreasing *N*. This nonlinearity implies a critical foot process number *N*_*c*_ below which the feedback reaches a mechanical point of no return, analogous to the critical crack length in fracture mechanics. The spatially ordered progression documented by Miyaki et al. [15], with edge loss preceding central loss, is consistent with this mechanism and difficult to explain by a diffusible toxin acting uniformly. From the translational perspective, the shear lag parameter *ω* could thus, in principle, serve as a patientspecific index of mechanical vulnerability, provided that the requisite morphometric inputs can be obtained from biopsy specimens via serial block-face scanning electron microscopy [15] or expansion microscopy [22]. Unlike proteinuria, which is downstream, *ω* reflects the structural mechanical state of the filtration barrier directly and might detect risk before functional decline becomes apparent. Therapeutically, the stiffness hierarchy suggests that cytoskeletal modulators targeting major process stiffness may relieve foot process overload more effectively than interventions aimed at GBM composition. The model also rationalises the renoprotective effects of SGLT2 inhibitors and RAS blockers, which reduce intraglomerular pressure [5] and thereby reduce the imposed displacement Δ. Because foot process force scales with Δ, even modest pressure reductions could meaningfully weaken the positive feedback loop. This mechanical interpretation complements the metabolic mechanisms conventionally invoked for SGLT2 inhibitor efficacy. These possibilities should be interpreted cautiously: the model predicts relative force distributions, not absolute magnitudes. Quantitative clinical predictions will require coupling to a haemodynamic model and validation against human data. The linear model also does not capture the viscoelastic, strain-stiffening response of the actin cytoskeleton [23, 24]. Nonlinear stiffening would amplify end-concentrated forces beyond the present predictions, suggesting that the present analysis provides a conservative estimate of force nonuniformity. The 2D geometry additionally neglects the curved nature of the capillary surface [7]. The model also represents the foot process-GBM attachment as a single linear spring (*k*_GBM_): because shear lag concentrates force at the segment ends, the integrin adhesions of the peripheral foot processes bear the largest load, a plausible physical trigger for the edge-first detachment seen in disease. A more complete model would resolve this attachment as a non-linear, force-dependent, remodelling adhesion (incorporating GBM viscoelasticity and disease-associated thickening) which could further strengthen the feedback loop. The disease application draws on morphometric data from a single animal model at three time points. Validation against genetic FSGS models, diabetic nephropathy, and human biopsy data constitutes an important future direction. Finally, the model does not incorporate mechanotransduction. Coupling the mechanical framework to a signalling model of cytoskeletal regulation, potentially closing the loop between force, TRPC6 activation [10], and structural remodelling and enabling prediction of how injury propagates.

## 5. Conclusions

This shear lag model of force transmission in the podocyte foot process network is, to our knowledge, the first analytical framework for predicting where mechanical failure initiates in the glomerular filtration barrier and how it propagates. The shear lag parameter *ω* distills foot process geometry, membrane stiffness, and cell mechanics into a single metric governing force decay. This could potentially serve as a structurally grounded biomarker of glomerular mechanical vulnerability, complementing functional readouts such as proteinuria. The model revealed that cytoskeletal stiffening of the major process amplifies foot process loading far more than equivalent changes in basement membrane properties, identifying a specific and potentially targetable mechanical driver of podocyte injury. Application to PAN nephrosis uncovered a positive feedback loop, in which force concentration drives foot process loss and thereby amplifies force on survivors. Results suggest a critical threshold below which mechanical failure becomes self-sustaining, regardless of the initiating insult. This threshold, if validated, could define the point at which podocyte injury becomes irreversible. More broadly, these results cast podocyte disease as a structural mechanics problem amenable to analytical prediction, and suggest a path toward patient-specific assessment of glomerular risk.

## Declarations

### Ethics

This study is purely computational and uses only previously published morphometric data from the literature. No new experimental work involving human participants or animals was conducted by the modelling team. Imaging data shown in Fig. 1 were generated under separate ethics approvals; please see the relevant Methods subsections.

### Data accessibility

The shear lag model is implemented in MATLAB; code and parameter files necessary to reproduce the results presented here are available at https://github.com/MonicaBBB/Shear-Lags.git.

### Author contributions

M.B.: conceptualisation, methodology, software, formal analysis, visualisation, writing: original draft; H.J.: conceptualisation, methodology, software, formal analysis, visualisation; P.P.: data curation, visualisation, investigation; Y.H.: methodology, software, formal analysis, visualisation; C.Q.: methodology, software, formal analysis, visualisation; J.H.M.: resources, methodology, supervision, funding acquisition, writing: review & editing; H.Y.S.: resources, methodology, investigation, funding acquisition, writing: review & editing; G.M.G.: conceptualisation, methodology, supervision, funding acquisition, writing: review & editing. All authors gave final approval for publication and agreed to be held accountable for the work performed therein.

